# Dynamic transcriptional remodeling in alcohol use disorder reveals immune dysregulation and adaptive shifts in coagulation during therapy

**DOI:** 10.64898/2026.05.15.725358

**Authors:** Susanne Edelmann, Thomas Hentrich, Sanya Esser, Sarah Pasche, Gilles Gasparoni, Mirac Nur Mosaoglu, Milan Zimmermann, Julia Schulze-Hentrich, Vanessa Nieratschker

**Affiliations:** Department of Psychiatry and Psychotherapy, Eberhard Karls University of Tuebingen, Tuebingen, Germany; German Center for Mental Health (DZPG), Partner Site Tuebingen, Germany; Department of Genetics/Epigenetics, Faculty NT, Saarland University, Saarbruecken, Germany; EpiGenomics Sequencing Facility, Saarland University, Saarbruecken, Germany

## Abstract

**Background:** Chronic alcohol use disorder (AUD) is associated with profound dysregulation of immune function, neuroinflammation, and systemic stress responses, which contribute to both the maintenance of addiction and alcohol-related organ damage. While brain transcriptomic studies have established neuroimmune signaling and synaptic remodeling as central features of AUD, peripheral blood signatures during early withdrawal and recovery remain underexplored. Understanding the dynamic transcriptional changes in peripheral blood accompanying supervised withdrawal therapy is critical for identifying reversible molecular processes versus persistent trait-like alterations.

**Methods:** RNA sequencing (RNA-seq) was performed on peripheral blood from individuals with alcohol use disorder (AUD, n = 100) and healthy controls (n = 74) at baseline and after three weeks of supervised withdrawal therapy. Differentially expressed genes (DEGs) were identified using linear mixed models assessing main effects of group, time, and their interaction. Functional enrichment and co-expression network analyses were performed to identify coordinated biological processes.

**Results:** At baseline, more than 1,000 genes were differentially expressed between AUD and control participants, showing robust dysregulation of immune-related pathways. After three weeks of withdrawal, the number of DEGs decreased markedly to 141, indicating partial transcriptomic normalization. Nevertheless, immune dysregulation persisted despite treatment, particularly linked to B cell activation and cell-cell junctions. Interaction analyses (group × time) identified 16 genes whose expression dynamically changed with therapy, highlighting strong enrichment for fatty acid pathways. Co-expression network analysis revealed that baseline modules were enriched for genes associated with secretory granules and immune signaling, while therapy-related co-expression shifts involved coagulation and platelet activation processes.

**Conclusions:** AUD is associated with widespread but partly reversible transcriptomic dysregulation in peripheral blood. These findings support a system-level view of AUD as a disorder of intertwined immune, metabolic, and coagulation biology and suggest that longitudinal blood transcriptomics may help identify both rapidly therapy-responsive and more stable molecular targets for relapse prevention.

## Introduction

Alcohol use disorder (AUD) represents a severe and multifactorial condition that imposes a major global health burden. According to the *World Health Organization Global Status Report 2024*, approximately 400 million individuals worldwide—corresponding to about 7% of adults aged 15 years and older—engage in harmful alcohol consumption. This behavior accounts for an estimated 4.7% of all global deaths (2.6 million annually) and contributes to 6.9% of male and 2.0% of female disability-adjusted life years (DALYs) in 2019 [1]. Despite its high prevalence and impact, effective long-term treatment options remain limited, with relapse rates exceeding 75% following initially successful interventions [2, 3]. The pathophysiology of AUD arises from the interplay between genetic predispositions and environmental factors, with gene–environment (G×E) interactions playing a pivotal role in determining individual vulnerability, therapeutic response, and relapse susceptibility [4–6].

Building on this framework of gene–environment interplay, large-scale RNA sequencing analyses in addiction-relevant brain regions, including the prefrontal cortex (PFC), nucleus accumbens (NAc) and amygdala, consistently implicate dysregulation of genes involved in inflammatory signaling, cell proliferation, and oxidative stress as core features of the AUD brain transcriptome: Recent work by Willis *et al*. systematically examined differential gene expression in the NAc and dorsolateral PFC from individuals with and without AUD, identifying hundreds of AUD-associated differentially expressed genes (DEGs) and region-specific co-expression networks [7]. Complementary to DEG-based approaches, Van Booven *et al*. demonstrated that AUD is accompanied by widespread alternative splicing abnormalities across multiple cortical and limbic regions [8]. Moreover, building on individual datasets, Friske and colleagues performed a systematic review and cross-species meta-analysis of 36 transcriptome-wide studies across PFC, NAc, and amygdala in humans, rodents, and non-human primates, revealing that the PFC harbors the largest burden of DEGs and that many of the most consistently altered genes cluster in neuroimmune pathways, including cytokine and interleukin signaling, *TNF*-related signaling, and microglial and astrocytic activation processes. [9]. Collectively, these studies indicate that chronic alcohol exposure induces pervasive changes in gene expression, with a strong and conserved enrichment of immune and inflammatory pathways in the addicted brain.

While brain tissue analyses provide direct insights into central pathophysiology, they are constrained by their post-mortem and cross-sectional nature, highlighting the need for translational approaches that capture dynamic, treatment-relevant molecular changes in living patients. Peripheral blood has therefore emerged as a complementary and clinically accessible compartment to probe alterations associated with alcohol exposure and AUD, and to translate brain-derived transcriptomic signatures into biomarker candidates that can be monitored longitudinally. Single-cell and bulk transcriptomic profiling of peripheral blood cells indicated that chronic alcohol use and heavy drinking were accompanied by pronounced remodeling of the immune system at the transcriptional level. Single-cell RNA sequencing of peripheral blood mononuclear cells revealed dose- and sex-dependent shifts in lymphocyte and monocyte proportions and cell type-specific changes in innate and adaptive immune gene expression, emphasizing complex immune dysregulation rather than isolated marker alterations [10]. Experimental primate models of chronic heavy ethanol consumption further demonstrated extensive differential gene expression in peripheral blood mononuclear cells, enriched for innate immunity, response to wounding, coagulation, and signaling processes, despite stable major immune cell subset frequencies, suggesting that alcohol reprograms immune cell function via transcriptional and stress-related mechanisms [11]. Integrative systems-biology frameworks proposed that such blood-based transcriptomic readouts, combined with other omics layers and advanced analytical approaches, can yield sensitive and specific RNA biomarkers to improve AUD diagnosis, patient stratification, and prediction of treatment response, by leveraging immune and oxidative stress–linked network signatures that reflect both central and peripheral disease processes [12].

Therapeutic interventions and withdrawal treatment add an important yet comparatively understudied dimension to this picture. Although most transcriptomic work in AUD has focused on cross-sectional post-mortem brain tissue or peripheral samples taken at a single time point, only a few studies have systematically examined how gene expression profiles change with treatment and how such changes relate to clinical response. One recent example is the small but informative study by Legaki *et al*., which specifically investigated neuroplasticity-related transcripts in the context of alcohol addiction and treatment. In that study, RNA from whole blood of 20 individuals with AUD and 10 healthy controls was analyzed before and after treatment using a targeted RT-PCR array comprising 84 neuroplasticity-related genes, identifying 35 genes as dysregulated in AUD and indicating broad alterations in synaptic signaling and plasticity pathways. Responders exhibited distinct baseline expression profiles, and post-treatment expression shifts toward normalization in a large subset of genes [13]. These findings provide first proof-of-concept that longitudinal peripheral blood gene expression profiling can capture treatment-related plasticity and may help stratify patients according to their likelihood of response, but they also underscore the need for larger, unbiased transcriptome-wide studies to robustly characterize molecular trajectories during withdrawal and early recovery in AUD.

Moreover, epigenetic mechanisms such as DNA methylation (DNAm), which act as environmentally responsive regulators of gene expression, have emerged as key molecular mediators of alcohol-related risk and recovery. Epigenome-wide longitudinal work in severely alcohol-dependent men has shown that acute withdrawal and short-term recovery are accompanied by extensive, partially reversible DNAm changes, with a strong over-representation of immune-related pathways, indicating that withdrawal and early abstinence are tightly coupled to dynamic immune epigenetic remodeling [14]. Similarly, DNAm profiling of purified CD3+ T cells in patients undergoing a three-week alcohol treatment program revealed widespread AUD-associated hypomethylation and a post-treatment shift of global and site-specific methylation patterns toward control levels, again highlighting immune cells as a critical compartment for alcohol-sensitive epigenetic regulation [15]. More recently, in our group, multimodal longitudinal analyses of the candidate gene *HECW2* across DNAm, gene expression, and histone marks in blood and saliva confirmed stable AUD-associated hypomethylation and therapy-related normalization of *HECW2* expression, thereby adding temporal and tissue-specific resolution to prior findings and underscoring the potential of DNAm and transcriptomic readouts as integrated biomarkers of AUD biology and treatment response [16].

However, longitudinal whole transcriptome studies in living patients, particularly in accessible tissues such as blood, are needed to identify transcriptional alterations appearing in AUD patients versus healthy controls and to determine in which biological mechanisms they are involved in and whether those alterations normalize with abstinence or persist as putative molecular scars associated with relapse risk and treatment outcome. In this context, peripheral transcriptomic profiling during supervised withdrawal and early recovery offers a translational view on the systemic adaptations in AUD and provides an opportunity to integrate findings with previous epigenomic studies and transcriptomic work in brain tissue to identify robust, treatment-relevant immune signatures.

The aim of this study was therefore to characterize longitudinal transcriptional changes associated with AUD and subsequent supervised withdrawal treatment. Using RNA sequencing of peripheral blood samples collected at baseline and after three weeks of inhouse qualified withdrawal treatment, we sought to identify differentially expressed genes and co-expression networks that reflect adaptive or persistent molecular alterations. Furthermore, we aimed to elucidate the biological pathways underlying early recovery from AUD and to evaluate whether transcriptomic signatures reproduce epigenetic patterns previously identified in epigenetic analyses, thereby providing a multi-layered perspective on the molecular mechanisms of treatment response and recovery.

## Methods

### Study population

In total, 174 participants of European ancestry (not genetically verified) aged between 22 and 71 years were included in the study. All patients suffering from AUD (n = 100) were diagnosed according to the International Statistical Classification of Diseases and Related Health Problems, 10th Revision (ICD-10, [17]) by experienced clinicians. Patients underwent a three-weeks inpatient withdrawal treatment at the Department of Psychiatry and Psychotherapy of the University Hospital Tuebingen. Samples and data were collected at hospital admission (T1), as well as after three weeks of therapy (T2). Accordingly, 74 healthy control individuals (HC) were also recruited twice, with a three-week interval between T1 and T2. At T2, 158 participants (N = 99 AUD patients, N = 59 HC) remained in the study. Of both groups, individuals with current substance use disorder other than nicotine or alcohol and with comorbid psychiatric disorders other than depression were excluded. At both time points alcohol consumption was assessed using the Alcohol Use Disorder Identification Test (AUDIT, [18]), and alcohol craving using the Obsessive-Compulsive Drinking Scale (OCDS, [19]) (Table S1).

All participants provided informed written consent. The study was approved by the ethics committee of the University of Tuebingen (Reference number 264/2018BO2 and 159/2021BO2) and was conducted in accordance with the Declaration of Helsinki.

### RNA extraction, library preparation and sequencing

Total RNA from whole blood stored in PAXgene Blood RNA tubes was extracted using the PaxGene Blood miRNA kit (Qiagen, Hilden, Germany). Quality of RNA was assessed using a Bioanalyzer (Agilent, Santa Clara, USA). Only samples with an RNA integrity number (RIN) of 7 and higher were used for sequencing library preparation. No sample was excluded at this point.

RNA-sequencing (RNA-seq) and library preparation were carried out in the Department of Genetics/Epigenetics, at the Saarland University, Saarbruecken, Germany, in two randomized batches. Sequencing libraries were prepared using the TruSeq Stranded mRNA Library Prep Kit (Illumina, San Diego, CA, USA). Sequencing was conducted on an Element AVITI system using Cloudbreak sequencing chemistry (AVITI 2×75 Sequencing Kit Cloudbreak FS High Throuput Kit, Element Biosciences, San Diego, CA, USA). For each sample, 35–50 million paired-end 2×75 bp reads were obtained.

### RNA-seq read preprocessing

RNA-seq read preprocessing and quality control were performed using the nf-core/rnaseq pipeline (version 3.19.0) with default parameters [20]. Raw FASTQ files were subjected to adapter trimming and quality filtering, followed by alignment to the human reference genome (GRCh38) using STAR+RSEM workflow (STAR for alignment, RSEM for gene/transcript quantification).

To account for heterogeneity in peripheral blood cell composition, cell type fractions were estimated from TPM expression values using CIBERSortX [21] with the LM22 leukocyte gene signature matrix, yielding estimated proportions for major immune cell subsets. Principal component analysis (PCA) of the CIBERSortX-derived cell type ratio matrix was performed using *prcomp* function (base stat package, version 4.4.3) with unit variance scaling, in order to derive orthogonal components summarizing major axes of variation in inferred cell composition; these principal components (PCs) were used as covariates in further analyses.

The contribution of technical and biological factors, including imputed cell fractions, to gene expression variability was quantified using the variancePartition package in R (version 1.36.3), by fitting linear mixed models to each gene using the following design: ~age + OCDS + AUDIT + (1∣group) + (1∣time_point) + (1∣ID) + batch + (1∣cohort) + (1∣sex) + (1∣smoking) + PC1 + PC2 + PC3 + PC4 + PC5.

The analysis indicated that imputed cell type composition explained only a minor fraction of expression variance, whereas substantial variance components were attributable to batch and other technical factors (Fig. S2), justifying explicit modeling of batch and additional random effects in downstream differential expression analyses. However, PC1, PC3 and PC5 were included in further analyses explaining the most variance of the principal components derived from cell fraction estimation.

### Data analysis

All statistical data analyses were performed using the software environments R and Python. All p values were corrected for multiple testing using the Benjamini-Hochberg method [22].

#### Demographic and Clinical Information

Normality of data was tested using Shapiro-Wilk test. The test revealed non-normal distributions for all variables (Table S2). Therefore, the comparison of the trait medians between the independent groups was performed using the Wilcoxon Mann-Whitney rank sum test. Associations between categorical variables were examined using chi-square tests. All functions were applied using the base stats package in R (version 4.4.3).

#### Differential Gene Expression (DGE)

Differential gene expression analysis was performed using DESeq2 (version 1.46.0 [39]), which analyzes differences in gene expression based on a negative binomial generalized linear model.

Housekeeping genes defined as highly expressed in >90% of controls and fold-change<1.2 (only 21) were removed as well as genes with low count defined as less than 10 counts in 10% of the samples. A linear model with the following formula was fitted: ~ group + time_point + group*time_point + age + sex + smoking + batch + PC1 + PC3 + PC5. The false discovery rate (FDR, to adjust the p value to multiple correction) was set as 0.05. Results were filtered for DEGs with an absolute log2 fold-change larger than 0.3.

#### Weighted Gene Co-expression Network Analysis (WGCNA)

Scale-free co-expression networks were constructed using the R package WGCNA that defines modules using a dynamic tree-cutting algorithm based on hierarchical clustering of expression values (minimum module size = 100, cutting height = 0.99). WGCNA was performed using filtered and variance stabilized count data (generated from the read count matrix using DESeq2’s getVarianceStabilizedData function).

The first network was constructed for T1 with a soft power of 10 at which the scale-free topology fit index approximated 0.9. To identify modules associated with therapy response, co-expression networks were constructed from subject-specific delta expression values (T2 minus T1), such that all network analyses were based on within-subject changes rather than absolute expression levels. For both, the batch correction function of sva::ComBat (version 3.54.0) was used [23]. The module eigenvalue was used to perform the Pearson correlation analysis (default) with the variables (i.e. questionnaire scores of OCDS and AUDIT as well as covariates age, sex, BMI, batch, all cell types and the principal components derived from the cell fraction estimation with each whole module. The interaction network of the WGCNA co-expression module “darkred” was visualized using Cytoscape (v3.10.3., [50]).

#### Gene Functional Enrichment Analysis

Gene list functional enrichment analysis was performed using the R package gProfiler2 (version 0.2.4 [43, 44]) by using the Gene Ontology (GO) resource (g:Profiler data release e113_eg59_p19_f6a03c19 for *Homo sapiens*, [45, 46]). Terms with FDR-corrected p values of <0.05 were considered significantly enriched.

#### Comparison with reference-based DNA methylation data

For the comparison with reference-based DNA methylation data, differentially expressed genes were intersected with genes associated with significantly differentially methylated cytosines reported in previous epigenetic studies of AUD. Specifically, we used CpG sites significantly associated with case–control status at T1 in whole blood from Witt *et al*. (supplementary “Table S2: CpG sites significantly associated with case control status at time point 1 after FDR correction”, [14], sites showing significant associations of alcohol use disorder with DNA methylation in the first blood replication cohort from Lohoff *et al*. (“Table S3: Overlap of probes from discovery and replication datasets showing significant associations of alcohol use disorder with DNA methylation” [24]. For the comparison of blood DNAm data associated with AUD, CpG sites significantly methylated in the context of disease status at T2 in whole blood from Witt *et al*. were used (supplementary “Table S3: CpG sites significantly associated with case control status at time point 2 after FDR correction”, [14].

## Results

### Demographic and clinical information

The study cohort included a total of 174 participants: 100 AUD patients and 74 healthy control individuals (Table 1, Table S1). First, cell type proportion estimation revealed widespread group differences with generally elevated immune cell proportions in patients (Fig. S1). Further, although age, sex, and smoking behavior of both groups were tried to be matched throughout the recruitment process, the groups revealed significant differences in those factors: AUD patients were in median 50 years old (median±IQR: 50±16.0), predominantly smoking (76%) and included more males (72%) than females (28%). Healthy control individuals were younger (43±23.0 years) and included less smokers (24%) as well as more females (50%) than AUD patients (p_age_=0.002, p_sex_=0.005, p_smoking_<0.001, Table 1). Alcohol Use Disorder Identification Test (AUDIT) scores of 26±10.4 at T1 and 23±13.2 at T2 were observed in AUD patients. As AUDIT scores of healthy control individuals were 3±3.0 at T1 and 3±2.0 at T2, the scores were significantly higher in AUD patients at both time points (p<0.001, Table 1). Obsessive-compulsive Drinking Scale (OCDS) scores were also higher in patients at both time points (AUD: 27±14.0 at T1 and 17±12.0 at T2; HC: 2±2.8 at T1 and 1±2.0 at T2, p<0.001, Table 1), which shows elevated craving and obsessive tendencies towards alcohol in AUD patients. Questionnaire scores improved post therapy in the AUD group, showing that the detoxification had positive effects on drinking behavior and psychological welfare of patients (Table S1).

**Table 1.** Demographics and alcohol related questionnaire scores of patients and healthy control individuals per timepoint. Either numbers (percentages) or median (±IQR) are given.

|  |  | AUD patients | Healthy control individuals | P-values |
| --- | --- | --- | --- | --- |
| Sample Size (N) |  | N = 100 | N = 74 |  |
| Sex | Female | 28 (28%) | 37 (50%) | $p=0.005$ |
|  | Male | 72 (72%) | 37 (50%) |  |
| Age (years) | | 50 (±16.0) | 43 (±23.0) | $p=0.002$ |
| Smokers | | 73 (76%) | 18 (24%) | $p<0.001$ |
| AUDIT | T1 | 26 (±10.4) | 3 (±3.0) | $p<0.001$ |
| | T2 | 23 (±13.2) | 3 (± 2.0) | $p<0.001$ |
| OCDS | T1 | 27 (±14.0) | 2 (±2.8) | $p<0.001$ |
| | T2 | 17 (±12.0) | 1 (±2.0) | $p<0.001$ |

### Differential gene expression in blood of AUD patients compared to healthy control individuals before and in response to therapy

Differential gene expression in blood of AUD patients compared to healthy controls before and in response to therapy showed a marked reduction in transcriptomic dysregulation over time. At baseline (T1), 1087 genes were significantly differentially expressed (FDR<0.05) between individuals with AUD and healthy controls (Fig. 1A,C, Table S3), indicating a broad disturbance of peripheral gene regulation.693 of the genes showed upregulation, whereas 394 genes were downregulated in AUD patients (Table S3).

**Figure 1.**
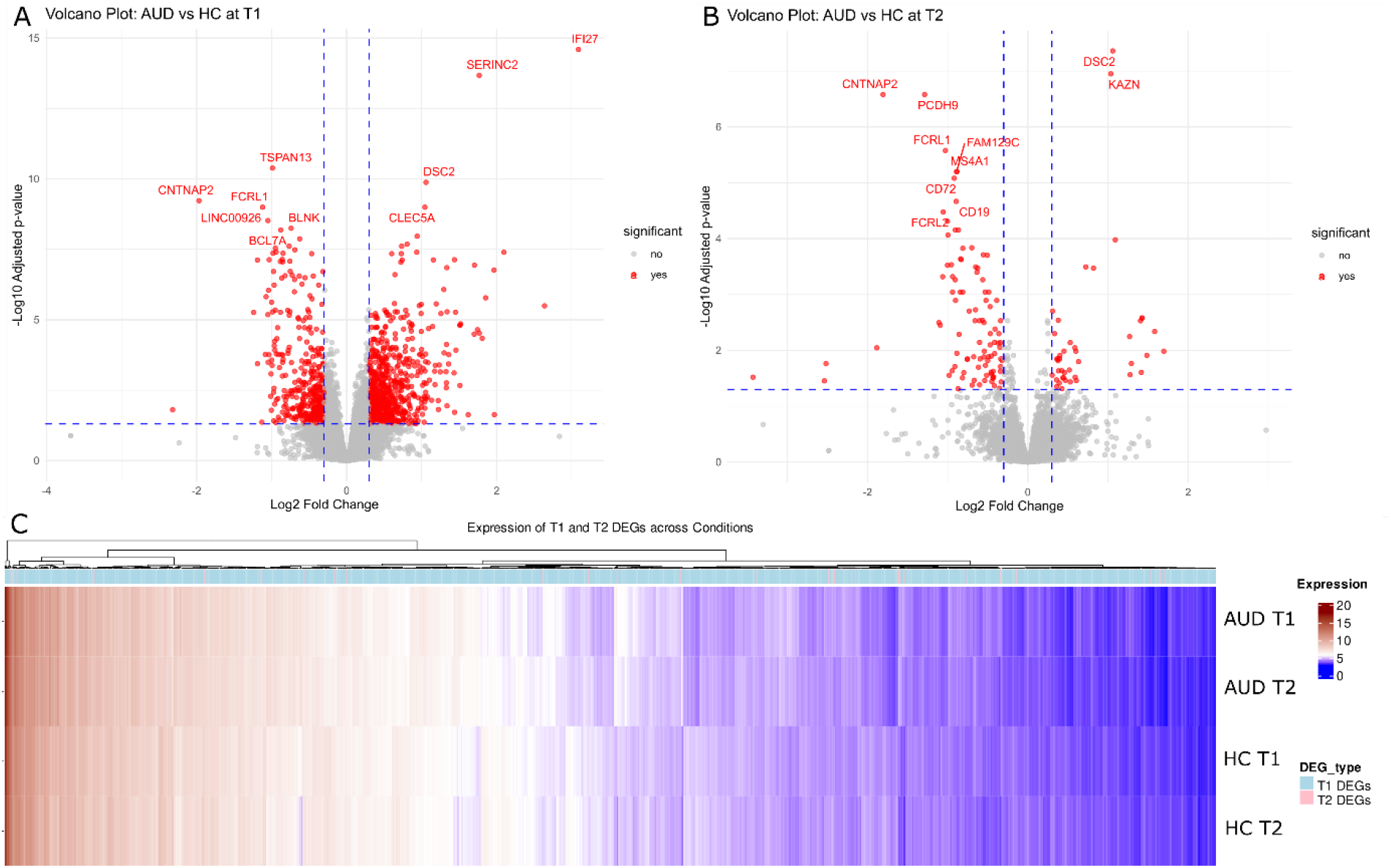
Differentially expressed genes at T1 (A) and T2 (B). Dashed lines indicate the statistical and fold-change thresholds applied; points are colored by significance. The ten most significant genes per comparison are labelled. **(C)** Mean expression (variance-stabilized and corrected for batch, PC1/3/5 as well as biological sex and age) of T1 and T2 DEGs across diagnostic group and time.

Functional enrichment analyses showed strong over-representation of broad immune system processes, defense responses, leukocyte activation, and genes annotated to the plasma membrane and cell periphery, consistent with pervasive activation and remodeling of circulating immune cells (Fig. 2).

**Figure 2.**
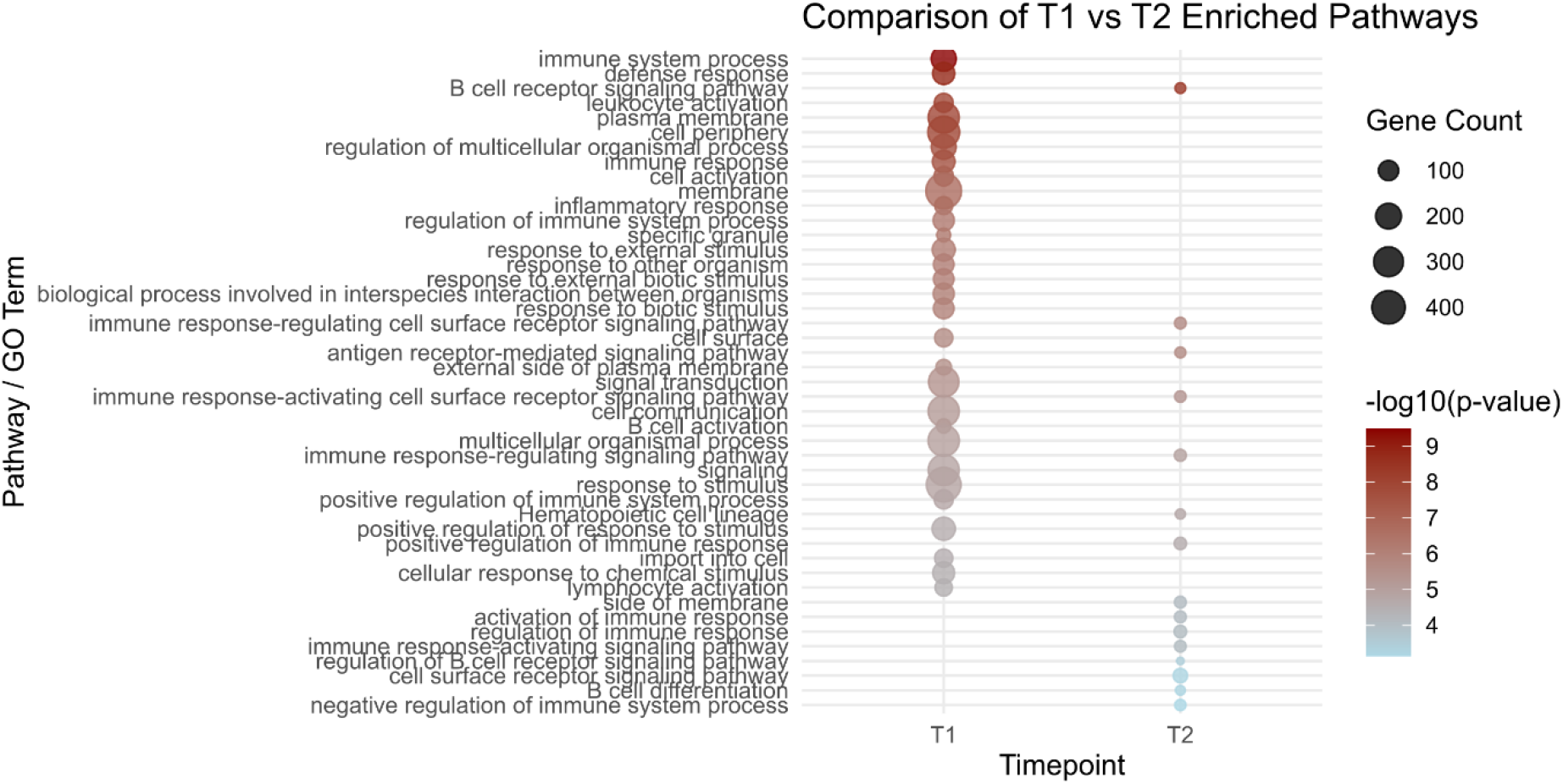
Functional enrichment analysis of the DEGs at T1 and T2. Top 40 enriched GO-terms are displayed, with bubble size indicating the number of genes per pathway and color representing enrichment significance (−log10 p value).

After three weeks of supervised withdrawal (T2), this signal was substantially attenuated, with only 141 genes remaining differentially expressed (FDR<0.05, with 98 down- and 43 upregulated, Fig. 1B, C, Table S4), highlighting B cell receptor signaling, immune response and immune system process terms, as well as cell-surface–localized genes and immune response–regulating receptor signaling pathways, suggesting a more restricted but still pronounced perturbation of adaptive and B cell–centered immune signaling under early abstinence (Fig. 2).

To specifically capture therapy-related dynamics, an interaction term (group × time) was modeled, identifying 16 genes whose expression trajectories over time differed significantly between AUD patients and controls (FDR<0.05, Fig. S3). Thirteen of the genes showed an increased difference over time in AUD patients compared to healthy controls (Table S5). *SERINC2* emerged as the top hit (interaction log2FC=−1.56, p<0.001), displaying higher expression in the AUD group across the observation period. Three genes revealed a decreased difference in expression over time in AUD patients compared to healthy controls (Table S5). Functional enrichment of these interaction-associated genes indicated involvement of fatty acid metabolic and biosynthetic processes, alongside terms related to cadherin-mediated cell–cell adhesion, cell–cell junctions, and adherent junctions. However, the enrichment signals were largely driven by only two to three genes, suggesting a limited but potentially biologically meaningful convergence of therapy-sensitive transcriptional changes on pathways integrating lipid metabolism with cell–cell contact and membrane organization during early recovery.

### Gene co-expression clusters associated with AUD and therapy

Co-expression analysis was performed to move beyond single-gene effects and identify sets of transcripts that change in a coordinated manner, thereby capturing higher-order pathway and cell-state signatures that may be more robustly linked to AUD severity and therapy response than individual DEGs.

At baseline (T1), 21 gene co-expression modules correlating with any of the variables available for the cohort (Table S1) were identified, with sizes ranging from 32 to 2128 genes. 5256 genes were assigned as not correlated (module grey). The darkred module including 32 genes was identified as the only co-expression cluster showing strong correlation with AUD severity (AUDIT: *r*=0.43, p<0.001; and OCDS: *r*=0.45, p<0.001, Fig. 3A, Fig. S4), while displaying only weaker associations with covariates such as age and inferred blood cell type proportions (BMI: *r*=0.11, p=0.159; age: *r*=0.16, p=0.042, PC1: *r*=0.19, p=0.015; PC3: *r*=0.09, p=0.258; PC5: *r*=−0.19, p=0.011, Fig. S3). Functional enrichment analysis at T1 revealed that this module was highly enriched for genes annotated to secretory granules, secretory vesicles, specific granules, and the lumina of secretory and cytoplasmic vesicles, indicating coordinated dysregulation of granule and vesicle biology in peripheral blood cells of individuals with AUD.

**Figure 3.**
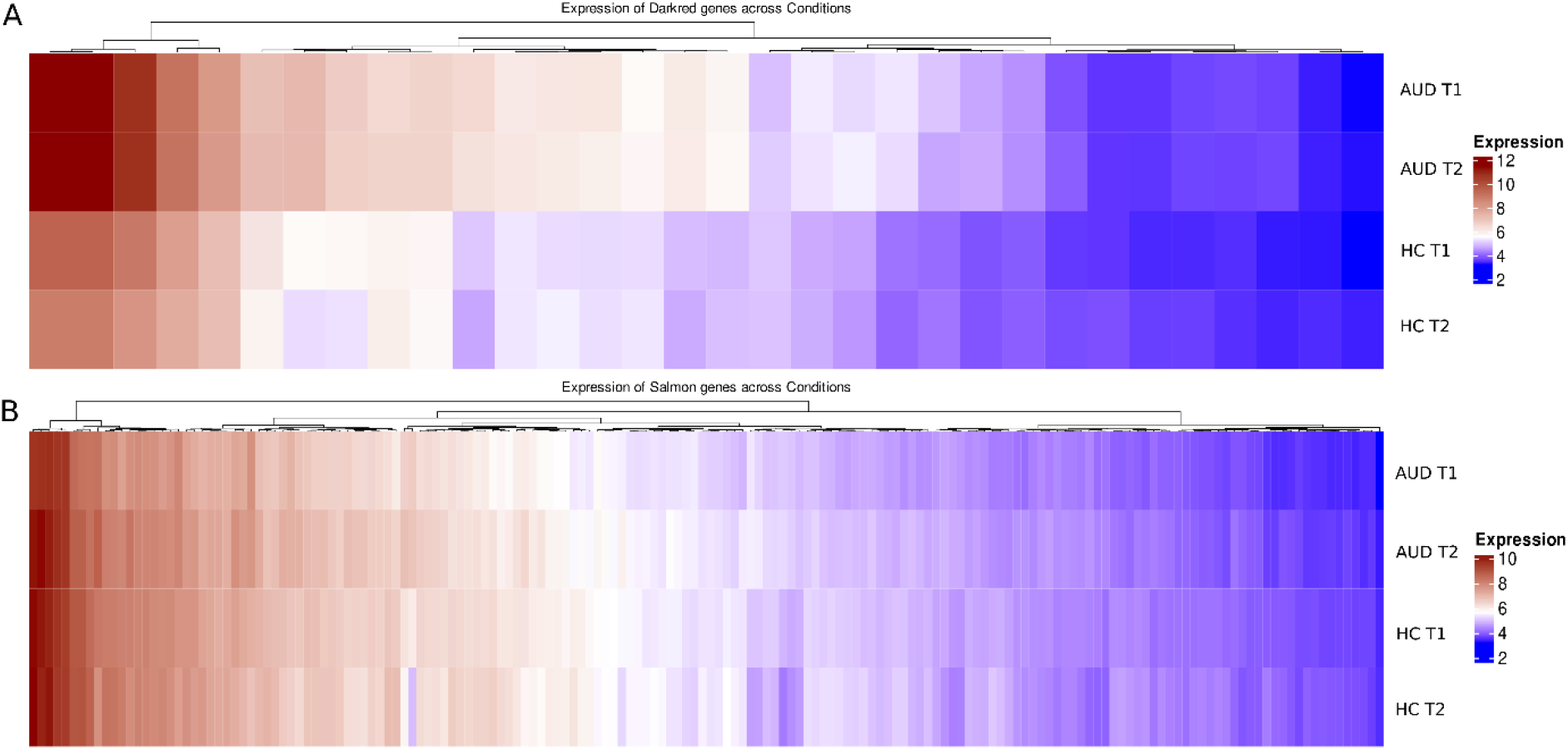
Mean expression (variance-stabilized and corrected for batch, PC1/3/5 as well as biological sex and age) of **(A)** genes included in the darkred co-expression cluster and **(B)** genes included in the salmon co-expression cluster across diagnostic group and time.

Among the genes of the darkred module, 25 were also assigned as baseline DEGs between AUD patients and healthy controls, forming a densely interconnected co-expression network, with genes showing high intramodular connectivity and clustering around a central core of strongly co-expressed nodes.

Co-expression clustering was also performed on within-subject expression changes (T2–T1) to identify modules associated with therapy response. The analysis revealed in total 20 clusters including 43 to 1988 genes. 5613 genes were assigned as not correlated with any variable (module grey). The salmon module including 169 genes emerged as the strongest therapy-responsive cluster (Fig. 3B), showing high treatment-sensitive correlations with AUD severity (AUDIT: *r*=0.29, p<0.001; and OCDS: *r*=0.23, p=0.003, Fig. S5), while correlations with covariates and principal components of inferred cell type ratios were comparatively modest (Fig. S5, BMI: *r*=0.03, p=0.704; age: *r*=0.09, p=0.266, PC1: *r*=0.06, p=0.441; PC3: *r*=−0.17, p=0.036; PC5: *r*=0.00, p=0.991), indicating that its signal was not primarily driven by demographic or compositional differences. Functional enrichment of the salmon module highlighted biological processes related to coagulation, hemostasis, and wound healing, as well as genes localized to platelet alpha granules (Fig. 4).

**Figure 4.**
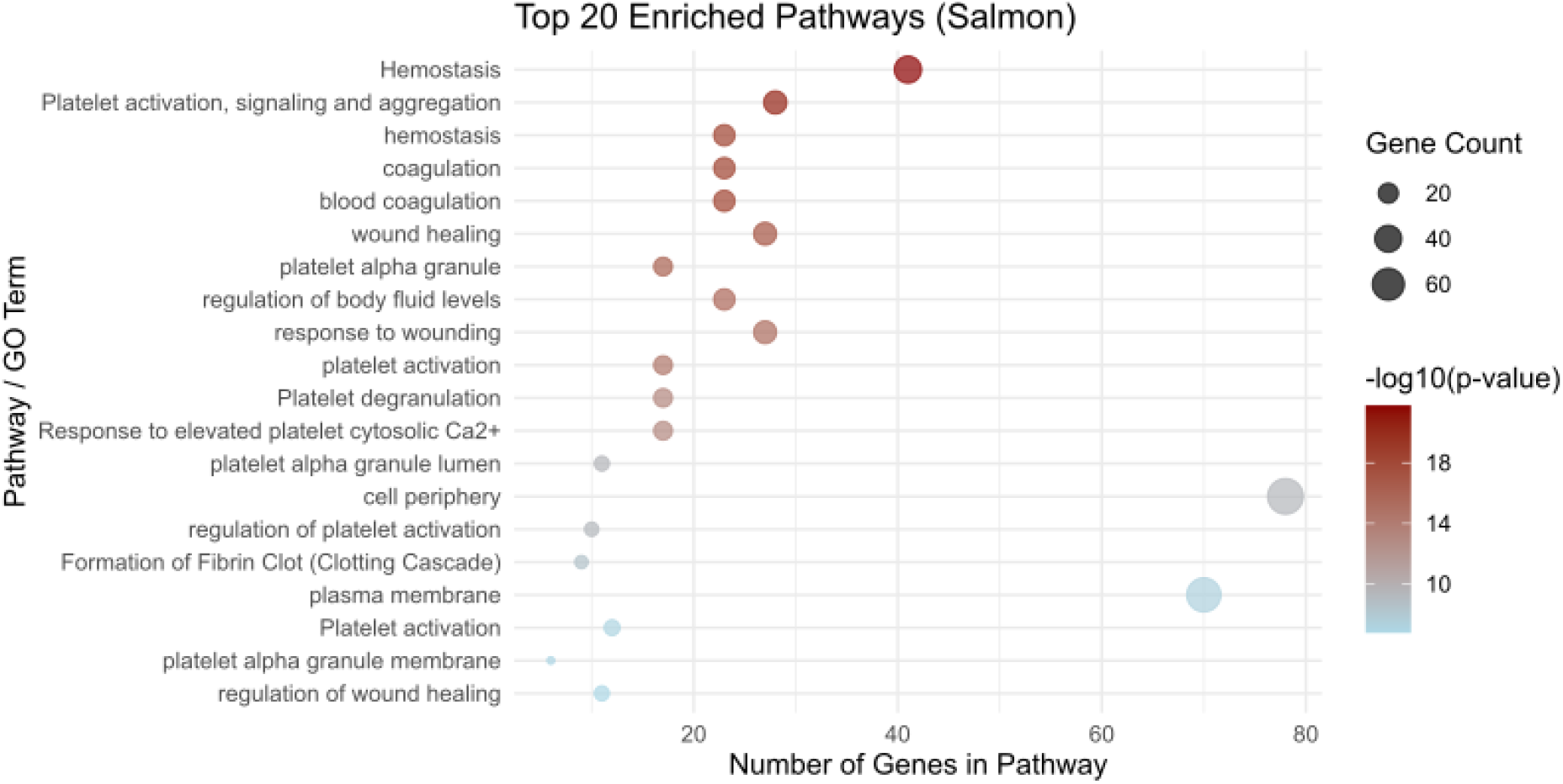
Functional enrichment analysis of the interaction-associated salmon co-expression cluster. Top 20 enriched GO-terms are displayed, with bubble size indicating the number of genes per pathway and color representing enrichment significance (−log10 p value).

Inspection of individual gene expression patterns within the salmon module showed that many of these coagulation- and wound-healing–related genes were upregulated in AUD patients at baseline (T1) compared with controls (Fig. S6A), whereas by T2 their expression differences were markedly attenuated (Fig. S6B).

### Epigenetic regulation of gene expression in AUD based on public data

Within a reference-based framework, genes highlighted by previous DNAm studies in AUD (CD3+ cells as well as whole blood) were intersected with the differentially expressed genes identified in this cohort, thereby restricting integration to loci that carry independent epigenetic evidence for alcohol-related regulation. The baseline transcription signature from the present sample showed partial overlap with whole-blood EWAS findings by Lohoff and colleagues, with three genes associated with differentially methylated cytosines ([24], Fig. 5). An EWAS conducted by Witt *et al*. in whole blood of men with alcohol withdrawal syndrome before and after therapy identified differentially methylated cytosines linked to numerous genes, of which 261 overlapped with the baseline DEGs detected in the present cohort, indicating convergent methylation–expression alterations at these loci under chronic alcohol exposure ([14], Fig. 5). One gene–*FKBP5*–was consistently shared across all baseline analyses.

**Figure 5.**
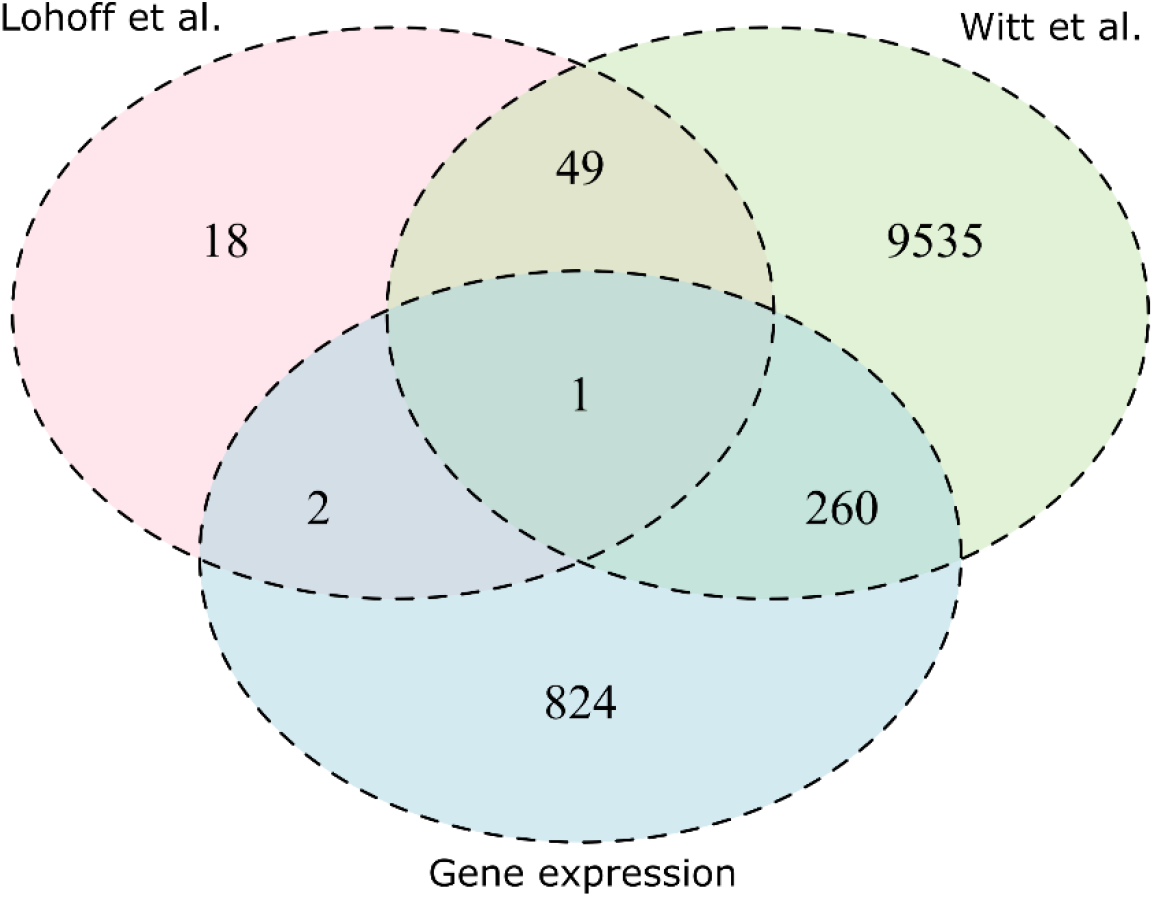
Overlap of genes across datasets and study types. Venn diagram illustrating the overlap of DEGs identified at T1 with genes associated with differentially methylated cytosines reported in previous DNA methylation studies by Lohoff *et al*. (pink) and Witt *et al*. (green), and with genes identified in this gene expression analysis (blue). Numbers indicate the count of genes unique to or shared between the respective datasets.

Intersection of genes associated with significantly differentially methylated cytosines following therapy (Witt *et al*.) and T2 DEGs was also conducted. The analysis revealed an overlap of 29 genes that are differentially expressed and associated with differential DNAm between AUD patients and healthy controls following a withdrawal treatment (Fig. S7).

## Discussion

In this study, we combined longitudinal gene expression analyses with functional enrichment and integrative comparisons to prior DNA methylation studies to investigate molecular processes associated with AUD and its treatment. We found significant and widespread changes in gene expression in the longitudinal comparison of AUD patients as well as between patients and controls. Interaction analyses identified a relatively small but specific subset of therapy-responsive genes, consistent with stricter statistical requirements for time-by-treatment effects. Co-expression network analysis revealed distinct gene modules linked to AUD status and treatment response, including clusters enriched for lipid- and coagulation-related pathways. Functional enrichment across these analyses highlighted coordinated dysregulation of immune and hemostatic processes, supporting a systems-level view of AUD involving intertwined inflammatory, metabolic, and coagulation-related pathways.

### Differential gene expression

The number of DEGs identified at T1 in the present study is broadly comparable to that reported in transcriptomic analyses of brain tissue in AUD, despite the clear differences in tissue context [9, 25]. Meta-analytic work by Friske and colleagues, for example, identified a set of common DEGs across brain regions, including *FKBP5*, with a predominance of upregulated signals in the prefrontal cortex and strong links to neuroimmune signaling pathways [9]. In our dataset, baseline DEGs in the context of AUD were also rather upregulated, even after accounting for changes in cell-type composition. This pattern supports a similar behaviors of gene expression patterns in blood and brain within the AUD phenotype, indicating shared molecular signatures across tissues [26]. At the same time, the only partial overlap in DEGs as reported by Friske *et al*. underscores pronounced tissue specificity, as peripheral immune cells and brain tissue differ markedly in their baseline regulatory programs, stress responsiveness, and epigenetic landscapes [9, 27]. However, *FKBP5* revealed to be consistently upregulated across different datasets and tissues as shown by Friske *et al*. and this study, supporting its potential as biomarker for AUD [9].

The present study demonstrated that individuals with AUD show robust alterations in immune and defense response pathways in peripheral blood at T1, supporting the view of AUD as a disorder with a pronounced (neuro)immune component [28]. These findings are highly consistent with prior work implicating immune and inflammatory mechanisms in AUD. Mayfield and colleagues have shown that chronic alcohol exposure alters expression of neuroimmune genes, including toll-like receptors, NF-κB-related components, and downstream cytokines, and that modulation of these pathways can influence alcohol-related behaviors in experimental models [29]. Crews and co-workers further highlighted that persistent activation of neuroimmune signaling contributes to alcohol-related neurodegeneration and cognitive impairment, placing inflammation centrally in AUD pathophysiology [30]. In line with this, the current blood-based results as well as epigenetic studies in peripheral tissue in the context of AUD [14, 24, 31–33] reinforce the concept that immune dysregulation is not only a correlate but a core molecular feature of AUD. The immune response pathways identified here at baseline may thus reflect a state of heightened systemic inflammatory tone that is sensitive to recent drinking and withdrawal status and potentially reversible with sustained abstinence. At the same time, the results need to be interpreted within the framework of neuroinflammation and tissue specificity. Neuroimmune models of AUD emphasize microglial activation, toll-like receptor signaling, and increased pro-inflammatory cytokines within specific brain regions such as prefrontal cortex and nucleus accumbens, where they influence synaptic plasticity, reward processing, and cognition [29, 34]. Peripheral blood, by contrast, primarily indexes systemic immune activity and may be more strongly shaped by circulating immune cells and peripheral cytokine milieu [35, 36]. Consequently, the immune response pathways detected in blood at T1 should be viewed as reflecting the systemic arm of the neuroimmune process rather than a one-to-one proxy for brain-region specific transcriptional modules. Beyond tissue specificity, our findings may also capture molecular processes that operate upstream or downstream of canonical signaling pathways such as *MAPK* or *STAT* signaling identified by Friske and colleagues. Peripheral transcriptional changes may reflect altered signaling thresholds, receptor availability, or metabolic constraints that modulate pathway activity observed in brain tissue [37]. In this framework, peripheral signals could represent upstream modulators of systemic stress and immune signaling, or downstream readouts of central or endocrine processes, rather than direct analogs of neuronal transcriptional responses.

To further contextualize our findings, we compared gene expression results with genes associated with differentially methylated cytosines reported in independent DNA methylation studies. The comparison of the baseline DEGs with existing epigenome-wide studies further supports an epigenetic contribution to immune-related dysregulation in AUD. Notably, the overlap of baseline DEGs with Witt’s differentially methylated cytosine-associated genes, where many alcohol- and withdrawal-associated DNAm changes map to immune and stress-related genes, is higher than with the EWAS conducted by Lohoff and colleagues, whose results directed to glucocorticoid and stress-system methylation signatures in large epidemiological cohorts [24]. The overlap of *FKBP5* across the publicly available DNA methylation datasets and our gene expression data appears both directionally and biologically plausible in the context of AUD and withdrawal. Lower methylation at *FKBP5* promoter CpGs (e.g. cg20813374 in Lohoff *et al*. and cg19226017 in Witt *et al*.) fits the canonical model in which promoter hypomethylation facilitates increased gene expression [38], which is in line with our results. However, intronic features often show more complex relationships between regulatory DNA methylation and gene expression than promoter CpGs, making a straightforward interpretation of directionality difficult [39, 40]. For the intronic CpGs identified by Witt and colleagues (cg03546163, cg14284211), the biological plausibility of the observed hypomethylation cannot be firmly established, but regulatory interactions—such as effects on intragenic enhancers or glucocorticoid-responsive elements—cannot be excluded.

A noteworthy feature of the longitudinal transcriptomic profile was the marked reduction in the number of DEGs from T1 to T2, which supports the idea that withdrawal treatment and early abstinence alleviate chronic cellular stress states and thereby permit partial re-regulation of gene expression. This pattern parallels longitudinal DNAm findings by Witt *et al*. and Brückmann *et al*., who observed extensive alcohol- and withdrawal-related DNAm changes that partially normalized after two to three weeks of therapy [15], particularly in genes involved in immune responses [14], suggesting that both epigenetic and transcriptional signatures are, at least in part, therapy-responsive. At the same time, a limited subset of genes remained dysregulated at T2, indicating that some transcriptional alterations may represent more persistent adaptations or be driven by regulatory mechanisms beyond acute transcriptional control, consistent with reports of residual epigenetic and immune-related changes after treatment and abstinence [14, 41]. Such residual dysregulation may point to molecular processes that are either less responsive to therapy or governed by regulatory mechanisms beyond acute transcriptional control. The functional profile of the remaining T2 DEGs, which were enriched for B cell–related and cell-surface–associated processes, fits with broader evidence that adaptive immune cell subsets and their surface receptors are altered in AUD and may recover only incompletely [42]. The B cell and membrane-related dysregulation could reflect long-lived immune memory or shifts in lymphocyte composition and is further reflected by the estimated cell type proportions in our data showing that naïve B cells were significantly lower in AUD patients compared to healthy controls at both time points.

To further shed light on the transcriptomic therapy response, we explicitly modeled the interaction between time and treatment, which highlighted a relatively small set of therapy-responsive DEGs—an expected finding given the stricter requirements of the interaction framework compared with simple T1–T2 comparisons, which capture all temporal changes irrespective of their specificity to therapeutic intervention [43]. Within this context, the identification of the top hit *SERINC2* as a therapy-responsive candidate is biologically plausible, given prior genetic work implicating *SERINC2* as a replicable risk gene for alcohol dependence and more recent evidence linking *SERINC2* variation to broader substance dependence and brain structural changes [44, 45]. Moreover, the involvement of genes related to cell– cell interactions and fatty acid or lipid metabolism is compatible with literature showing that AUD perturbs membrane composition, lipid handling, and immune cell function [46, 47], underscoring that the observed transcriptomic therapy response likely reflects coordinated adjustments in both immune and metabolic pathways rather than isolated gene-level effects. These observations suggest that while some AUD-associated transcriptional alterations normalize over time, others persist in a tissue-specific and pathway-dependent manner, underscoring the need for systems-level approaches such as co-expression network analysis to disentangle coordinated, cluster-level adaptations from isolated gene-level effects.

### Co-expression analyses

At T1, the co-expression and functional analyses highlighted a module (darkred) that was strongly associated with dysregulation of genes involved in secretory granules and secretory vesicles, indicating disruption of granule and vesicle biology in peripheral blood cells of individuals with AUD prior to therapy. This module showed substantial overlap with T1 DEGs and was centered on *CEACAM8* as a hub gene, a granulocyte-lineage activation marker stored in neutrophil secondary granules and secretory vesicles that is rapidly mobilized upon stimulation and participates in cell adhesion, degranulation, and immunoregulatory signaling [48]. Together with the immune response enrichment observed at T1, these findings suggest that pretreatment, blood of AUD patients is characterized at the transcriptomic level by a coordinated dysregulation of neutrophil/granulocyte secretory granule pathways and vesicle-mediated effector functions, consistent with a state of chronic immune activation and cellular stress [49, 50] that likely reflects more than direct oxidative toxicity of ethanol alone and instead points to broader remodeling of innate immune effector machinery.

The network structure supports the notion that AUD at T1 is characterized by coordinated perturbation of secretory granule that affect the immune response. Secretory granule pathways play a central role in innate immunity by controlling how neutrophils and other granulocytes store and rapidly release effector molecules such as proteases, antimicrobial peptides, and adhesion receptors [51]. Upon activation, granule and vesicle contents are mobilized to the cell surface or extracellular space, enabling fast antimicrobial killing, endothelial adhesion, tissue migration, and modulation of inflammatory signaling [52]. Consequently, dysregulation of secretory granule biology in AUD blood at T1 suggests an altered readiness and quality of neutrophil effector responses, compatible with a chronically activated yet potentially dysbalanced inflammatory state.

In the longitudinal interaction analysis, the salmon co-expression module emerged as correlated with therapy-responsive measures of AUD severity (AUDIT and OCDS) and was enriched for coagulation-related processes, in line with clinical evidence that heavy alcohol use alters hemostatic balance and can promote a procoagulant state through effects on platelets, coagulation factors, and inflammation-driven activation of the coagulation cascade [53, 54]. Within this module, overlap with interaction DEGs implicated genes involved in fatty acid and lipid metabolic processes, which is biologically plausible because fatty acid–dependent membrane composition shapes endothelial and immune cell membranes, stabilizes intercellular junctions [55], and thereby influences vascular barrier integrity and exposure of procoagulant surfaces that regulate coagulation [56].

These systems-level patterns support a model in which AUD-related transcriptomic dysregulation before therapy reflects not only direct ethanol toxicity but a broader, stress-amplified disturbance of mitochondrial, lipid, immune, and coagulation biology. Chronic alcohol exposure is associated with persistent oxidative stress and elevated reactive oxygen species (ROS, [57]), which damage mitochondrial function and deplete redox cofactors such as NADPH and glutathione [58]. This in turn constrains fatty acid biosynthesis, lipid remodeling, and antioxidant regeneration, shifting metabolism from an adaptive to a reactive mode [59]. At the membrane level, ROS-driven lipid peroxidation and impaired lipid turnover can compromise membrane composition and organization, contributing to mislocalization of junctional proteins, receptors, and coagulation regulators and thereby promoting chronic junctional leakiness and endothelial dysfunction [60]. Within this framework, dysregulated fatty acid metabolism, membrane biology, and secretory granule pathways in AUD blood before therapy can be viewed as interconnections of a low-energy, high-ROS state. Secretory granule function is tightly dependent on intact membranes, cytoskeletal dynamics, calcium handling, and ATP availability [61]. Therefore, energy and calcium stress, together with structural instability, can lead to defective granule trafficking, premature or incomplete priming, and inappropriate release of vWF, P-selectin, ADP, serotonin, and inflammatory mediators [62]. Loss of spatial and temporal control over secretion, in turn, lowers the activation threshold of platelets and leukocytes [63] and feeds into a procoagulant surface phenotype characterized by persistent phosphatidylserine exposure and increased tissue factor accessibility on activated platelets and endothelium, providing catalytic platforms for tenase and prothrombinase complex assembly [64, 65]. Over time, this combination of uncontrolled secretion and procoagulant surface dominance may favor chronic coagulation and immunothrombosis [66], with sustained thrombin generation, microclot formation, platelet–leukocyte aggregates, and local hypoxia, which further enhances ROS production and establishes a feed-forward, self-stabilizing state of low energy, high inflammation, and heightened thrombotic risk.

The longitudinal data suggest that withdrawal treatment interrupts this cycle to a considerable degree. Reducing or eliminating ethanol intake is expected to lower ROS levels and relieve ongoing oxidative pressure, which can rapidly stabilize lipids and membranes even if deeper structural and mitochondrial damage persists [67]. Fatty acid metabolism and coagulation are particularly ROS-sensitive bottlenecks: Enzymes of fatty acid biosynthesis and desaturation, as well as polyunsaturated lipid species, are directly vulnerable to oxidative modification [68], while coagulation depends on membrane lipid asymmetry, phosphatidylserine exposure, and oxidized lipid signaling at platelet and endothelial surfaces [69]. Consequently, modest reductions in ROS can produce functional gains in these pathways—less lipid peroxidation restores key metabolic enzyme activity and membrane lipid composition, and small shifts in phosphatidylserine exposure, tissue factor activation, and platelet reactivity can move the coagulation system to a more controlled state. In this sense, improvements in fatty acid metabolism and coagulation after therapy likely reflect the chemical reversibility and threshold behavior of these systems, rather than complete resolution of underlying cellular injury. By contrast, the persistence of T2 dysregulation in cell-surface organization and B cell–related processes indicate domains that are less rapidly reversible by redox normalization. Surface protein topology, receptor composition, and junctional architecture are shaped by structural, developmental, and trafficking programs that tend to be “hard-wired” [70, 71] and may require active reprogramming rather than simple removal of oxidative insults. Similarly, B cell activity depends on prior differentiation history, germinal center selection, and epigenetic imprinting of transcriptional programs [72]. Chronic oxidative and inflammatory stress during activation can leave an “immune memory” in terms of altered thresholds for activation, tolerance, or exhaustion, which is not automatically erased when ROS levels fall [73]. Consistent with this, the remaining T2 DEGs were enriched for B cell and cell-surface– associated pathways, and estimated cell-type proportions showed significantly reduced naïve B cells in AUD patients compared with controls at both time points, suggesting that adaptive immune remodeling persists beyond the acute detoxification window.

Overall, these observations support a layered model in which AUD-related oxidative stress, mitochondrial strain, and lipid dysregulation propagate outward through membranes, junctions, granules, and coagulation to generate a proinflammatory, prothrombotic milieu, while withdrawal therapy preferentially normalizes those components that are chemically and functionally most reversible—particularly fatty acid metabolism and coagulation—yet leaves more deeply embedded, structurally and epigenetically encoded alterations in cell-surface organization and B cell function only partially resolved within the studied timeframe. Current evidence suggests that detoxification mainly targets acute withdrawal physiology, while some of the molecular changes discussed above likely outlast the detox period and may contribute to relapse vulnerability. Standard medical treatment of AUD focuses on symptom-triggered benzodiazepine regimens, thiamine supplementation, and supportive care [74, 75], but does not directly address persistent neuroimmune, metabolic, or epigenetic alterations. Beyond pharmacological therapy, guideline-based treatment now routinely integrates psychosocial interventions, stress-management strategies, and relapse-prevention pharmacotherapy (e.g. naltrexone and acamprosate), which indirectly target brain circuits involved in stress responsivity, craving, and habit formation [75, 76]. Work on stress systems and FKBP5 illustrates how brain-level mechanisms might link molecular “scars” to relapse risk. Variants and expression changes in *FKBP5* have been associated with withdrawal severity [77], HPA axis dysregulation [78], and stress-related relapse vulnerability in AUD [79], and experimental studies indicate that *FKBP5* modulation can influence alcohol-related behaviors and stress reactivity [80]. More broadly, preclinical and human data show that alcohol-related epigenetic and neuroimmune changes in brain may persist into abstinence and are implicated in anxiety, cognitive inflexibility, and heightened relapse risk, although the exact duration and reversibility likely vary across individuals and brain regions [81–83]. These remaining questions—how best to support detoxification at the molecular level, how long structural and epigenetic damage persists, and to what extent such changes enhance relapse risk— underscore the need for integrated approaches that combine medical withdrawal management with targeted interventions on stress circuitry, neuroimmune signaling, and long-term behavioral support.

## Conclusion

These findings indicate that AUD is associated with widespread but partly reversible transcriptomic dysregulation in blood, with pronounced involvement of immune, metabolic, and coagulation-related pathways. Longitudinal patterns suggest that withdrawal treatment and abstinence can normalize stress- and ROS-sensitive processes, particularly fatty acid metabolism and hemostatic pathways, whereas alterations in cell-surface organization and B cell–related programs appear more persistent and may contribute to longer-term relapse vulnerability. Together, these results argue for a layered model in which epigenetically regulated peripheral transcriptomic markers capture both rapidly therapy-responsive and more stable, trait-like components of AUD biology, underscoring the value of integrating longitudinal blood-based omics with brain-focused, epigenetic, and clinical data to refine future biomarker development and intervention strategies.

## Limitations

Several limitations should be considered when interpreting these findings. The AUD and control groups were not fully balanced with respect to age, sex, and smoking status, all of which are known to exert substantial effects on blood gene expression and epigenetic profiles, particularly in immune and inflammatory pathways. However, these variables were included as covariates in the statistical models, which should mitigate—but cannot entirely eliminate—their potential confounding influence. Furthermore, although cell type deconvolution was applied, and cell type ratios were controlled for in further analyses, the observed differences in estimated blood cell composition between AUD patients and healthy controls imply that residual confounding by cell-type abundance may persist, particularly for rare cell populations and for signals that are sensitive to modeling assumptions underlying the deconvolution approach. Moreover, the study focused on mRNA and reference DNA methylation signatures and did not assess other regulatory layers such as microRNAs, which are known to modulate alcohol-related gene networks in brain and blood and can mediate regulatory miRNA changes in AUD [84].

## Outlook

Future work should build on the multi-omics approaches using longitudinal frameworks with larger sample sizes to identify treatment-responsive factors and validate diagnosis-associated factors, complemented by drug repositioning analyses to identify novel therapeutic candidates.

In parallel, more detailed *FKBP5* analyses across blood and brain-focused datasets could help clarify how stress-regulatory circuitry and HPA-axis related epigenetic marks contribute to withdrawal severity and relapse risk.

In addition, targeted assessment of mitochondrial DNA (mtDNA) damage and copy number, given evidence that chronic alcohol exposure induces mtDNA damage and degradation, could quantify the extent of mitochondrial involvement in the low-energy, high-ROS state inferred from the present data. Extending beyond mRNA and DNAm to other omics layers, including miRNAs and other non-coding RNAs, will be important, as recent work suggests that alcohol-induced miRNA changes can reprogram metabolism and neuronal signaling and are emerging as promising biomarkers and therapeutic targets [85].

Finally, longer-term follow-up combining molecular profiling with detailed clinical characterization of stress, comorbid psychopathology, and relapse outcomes will be required to test the hypothesis that reducing ROS and acute cellular stress, while necessary, is not sufficient to reverse deeper structural and epigenetic changes, and to evaluate whether persistent alterations in immune, metabolic, and stress-regulatory pathways indeed mark individuals at elevated risk for relapse.

## Author Contributions

SE: conceptualization, funding acquisition, project administration, resources, investigation, formal analysis, methodology, visualization, supervision, writing - original draft, writing – review & editing; TH: formal analysis, methodology, writing – review & editing; SE: methodology, writing – review & editing; SP: methodology, writing – review & editing; MZ: methodology, writing – review & editing; JSH: resources, writing – review & editing; VN: conceptualization, funding acquisition, project administration, resources, supervision, writing – review & editing

## Institutional Review Board Statement

This study was conducted in accordance with the Declaration of Helsinki and approved by the Institutional Ethics Committee of the University of Tuebingen (159/2021BO2 as well as 264/2018BO2). Informed consent was obtained from all participants involved in the study.

## Data Availability Statement

The original contributions presented in this study will be published with European Genome Archive (https://ega-archive.org/). Further inquiries can be directed to the corresponding author.

## Acknowledgements

We would like to express our appreciation to all participants for their contributions. Furthermore, we thank all residents present during the time of recruitment for contributing to the recruitment process. Moreover, we thank the Epigenomics Sequencing Facility of the Saarland University for library preparation, quality control and sequencing of the samples. We furthermore thank the DZPG for infrastructural support.

