## Supplemental Figures for "Dynamic transcriptional remodeling in alcohol use disorder reveals immune dysregulation and adaptive shifts in coagulation during therapy"

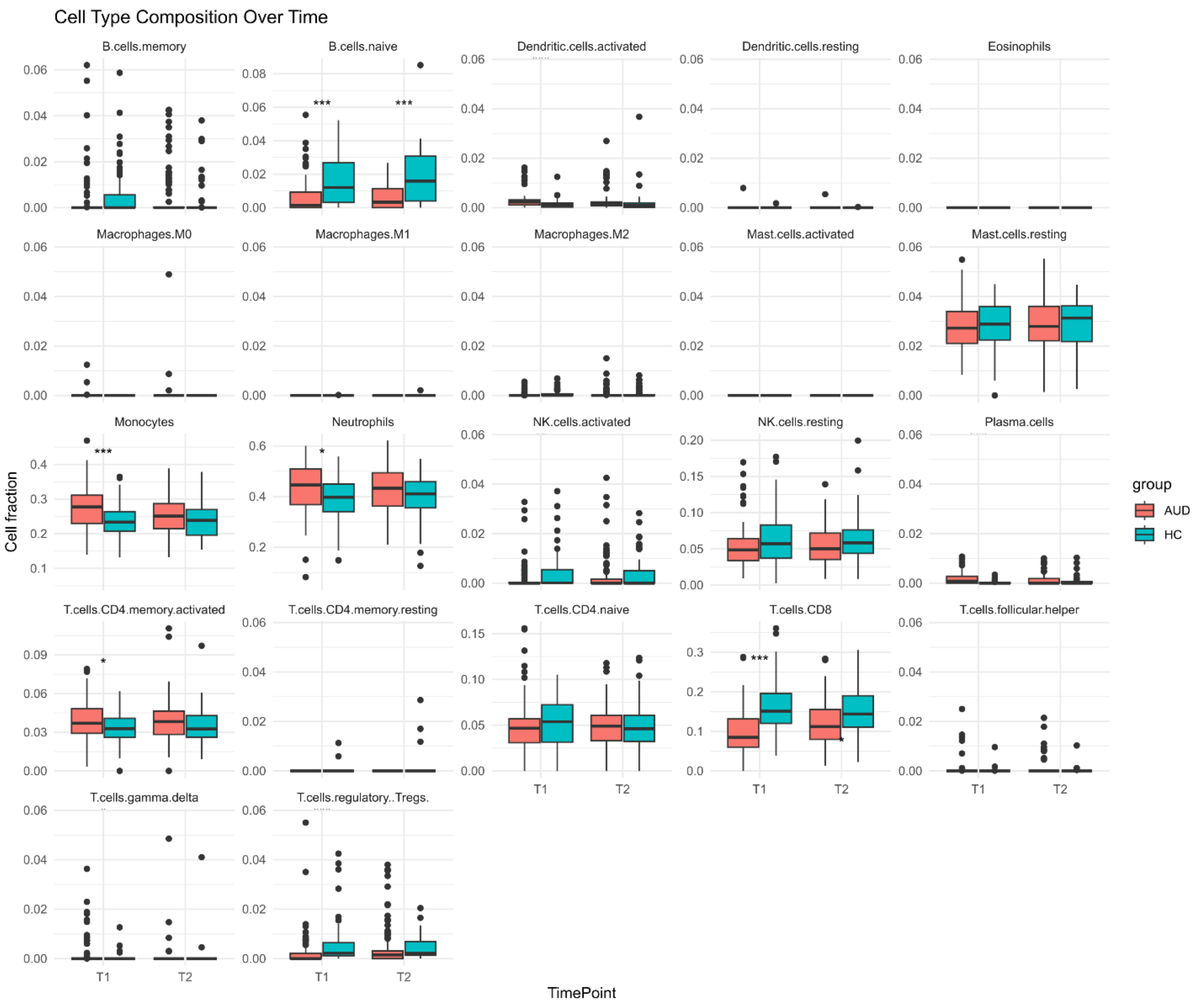


**Figure S1**: Estimated cell type proportions from blood of AUD patients and healthy control individuals divided by time point.

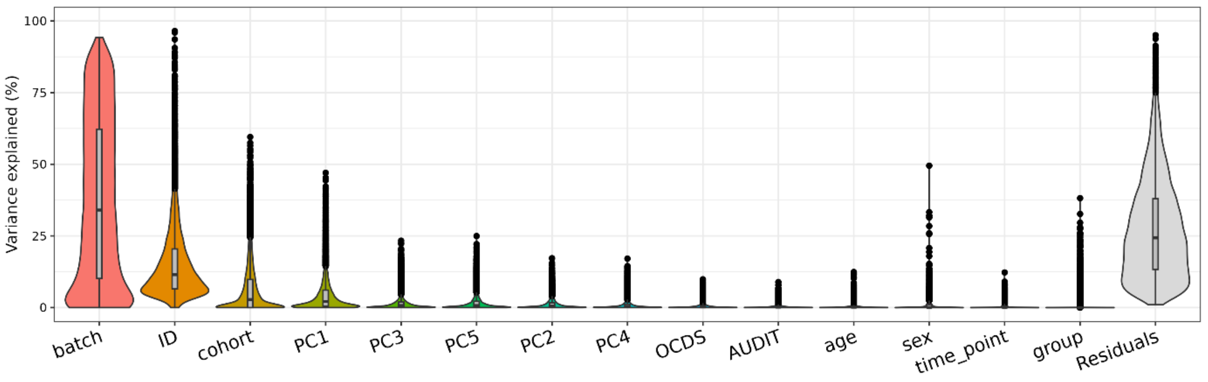


**Figure S2**: Variance partitioning of gene expression.

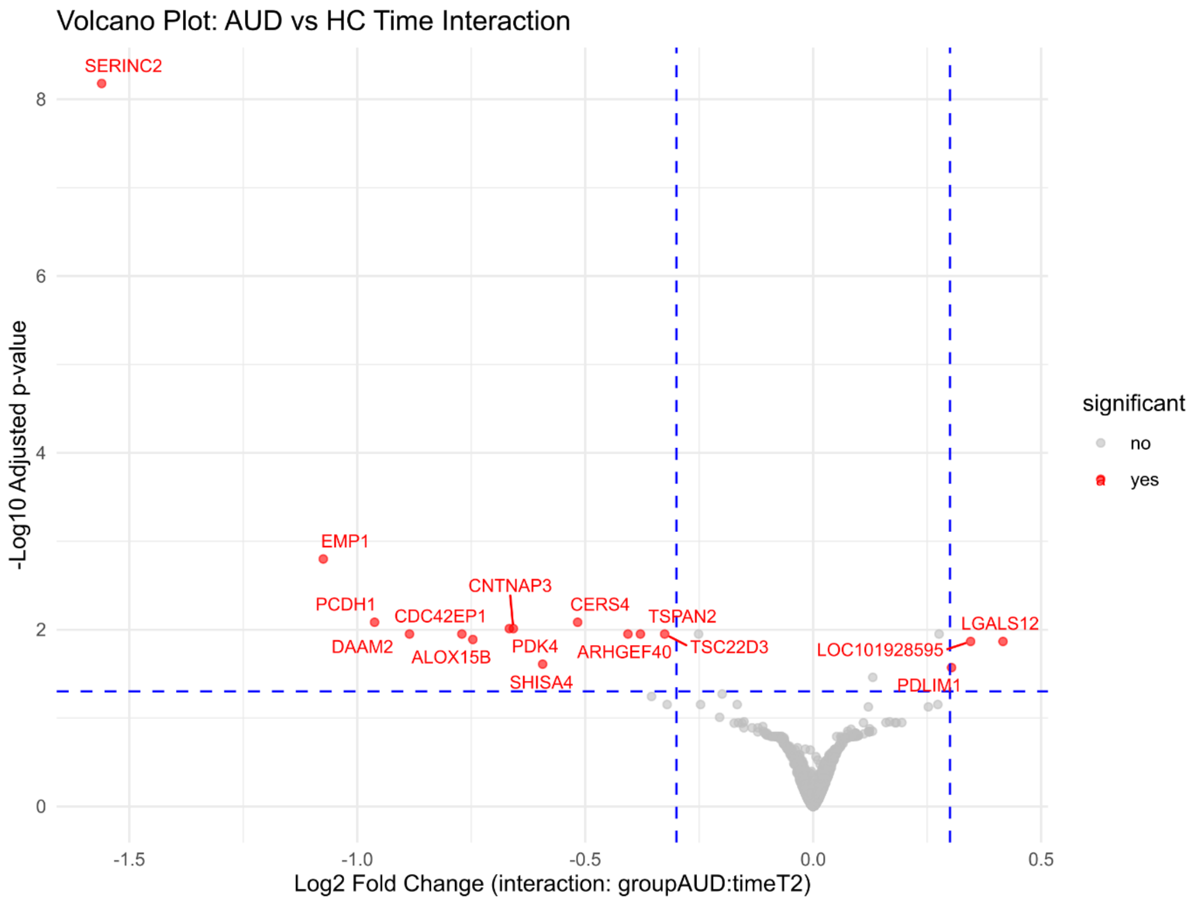


**Figure S3.** Differentially expressed genes of the interaction term (group × time), where expression trajectories over time differing significantly between AUD patients and controls were modelled. Dashed lines indicate the statistical and fold-change thresholds applied; points are colored by significance. All significant genes are labelled.

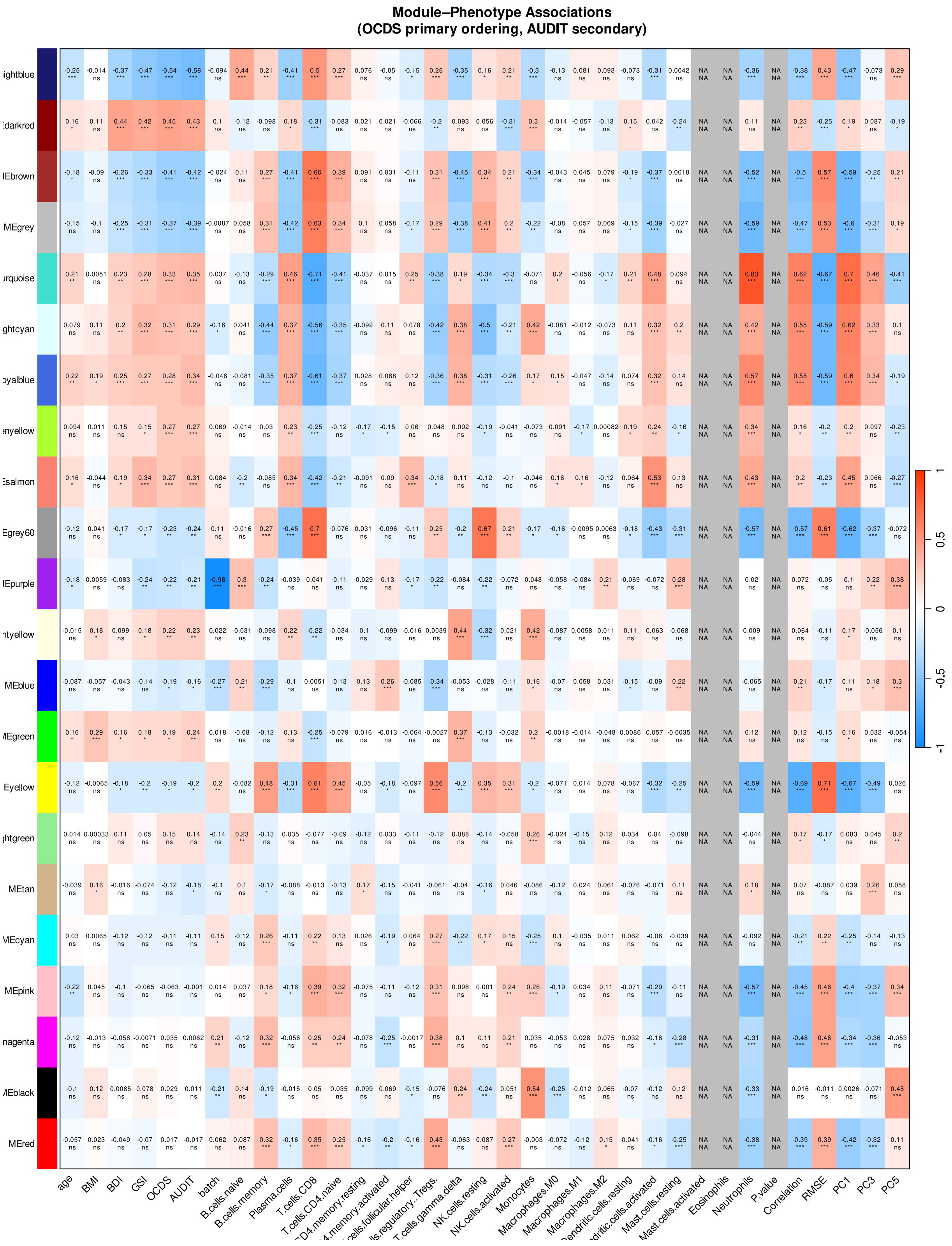


**Figure S4:** Module-Phenotype correlations of the darkred module (co-expression cluster T1)

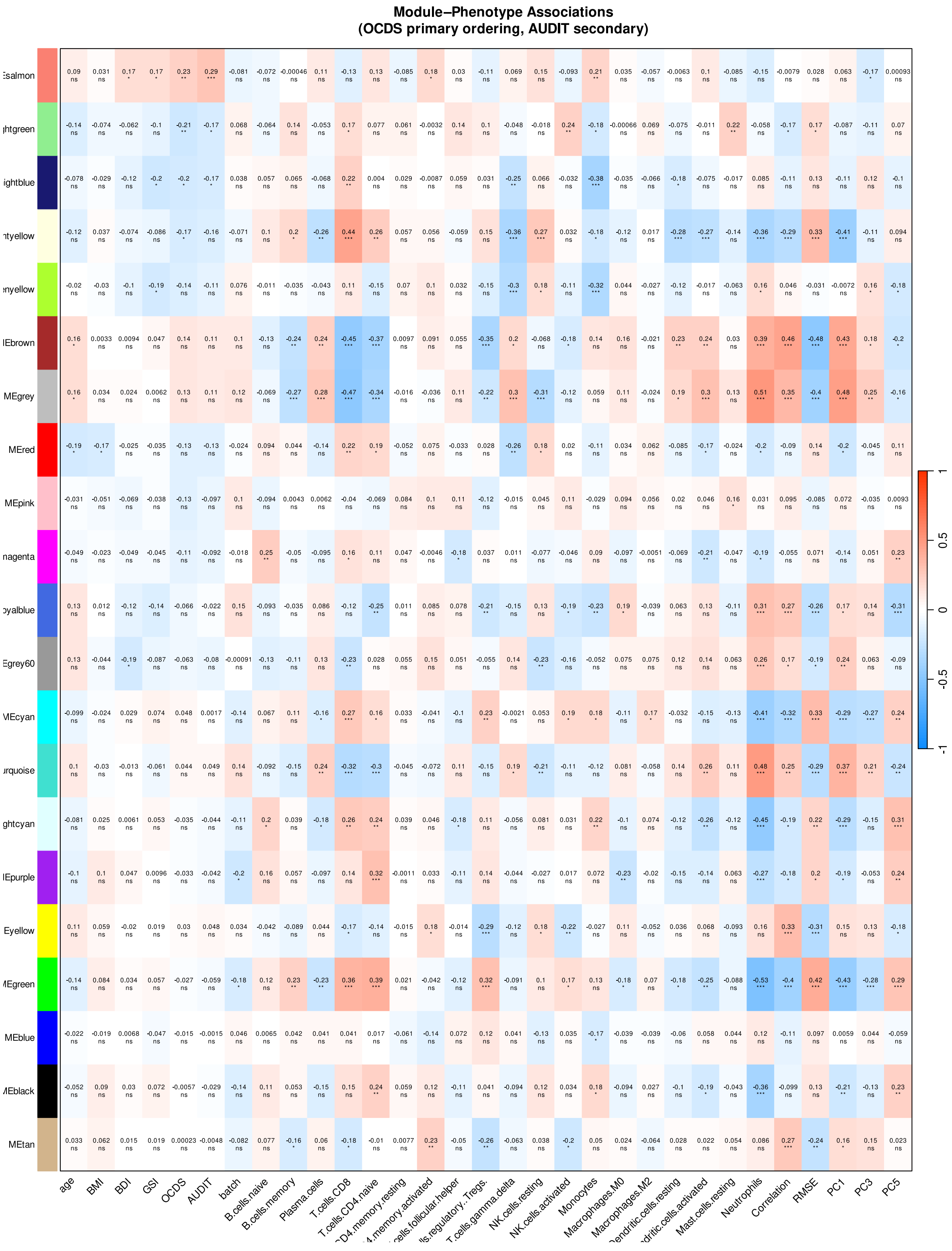


**Figure S5:** Module-Phenotype correlations of the salmon module (co-expression cluster T1-T2)

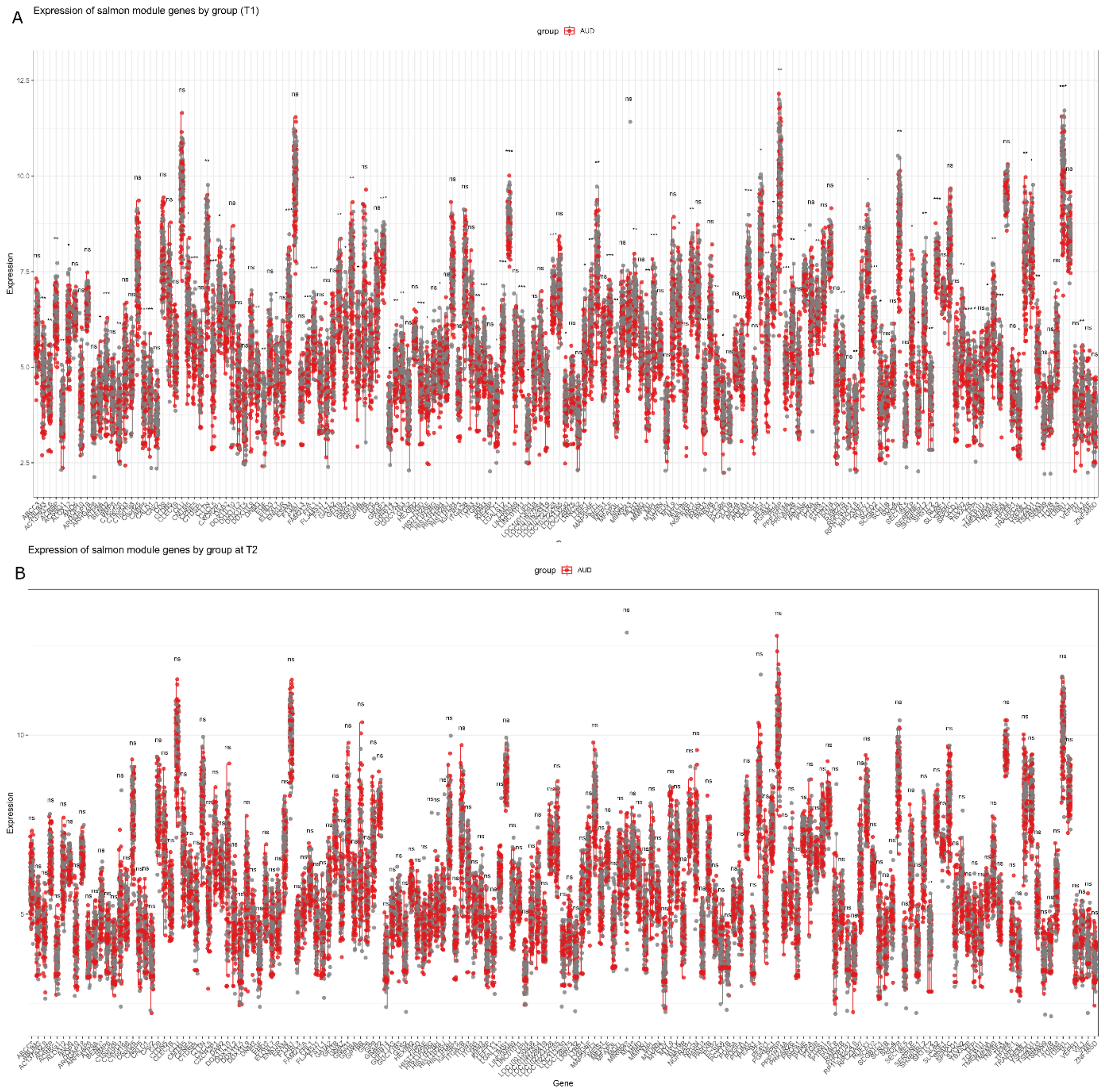


**Figure S6:** Expression of genes included in the salmon module (co-expression cluster T1-T2) at T1 (A) and T2 (B)

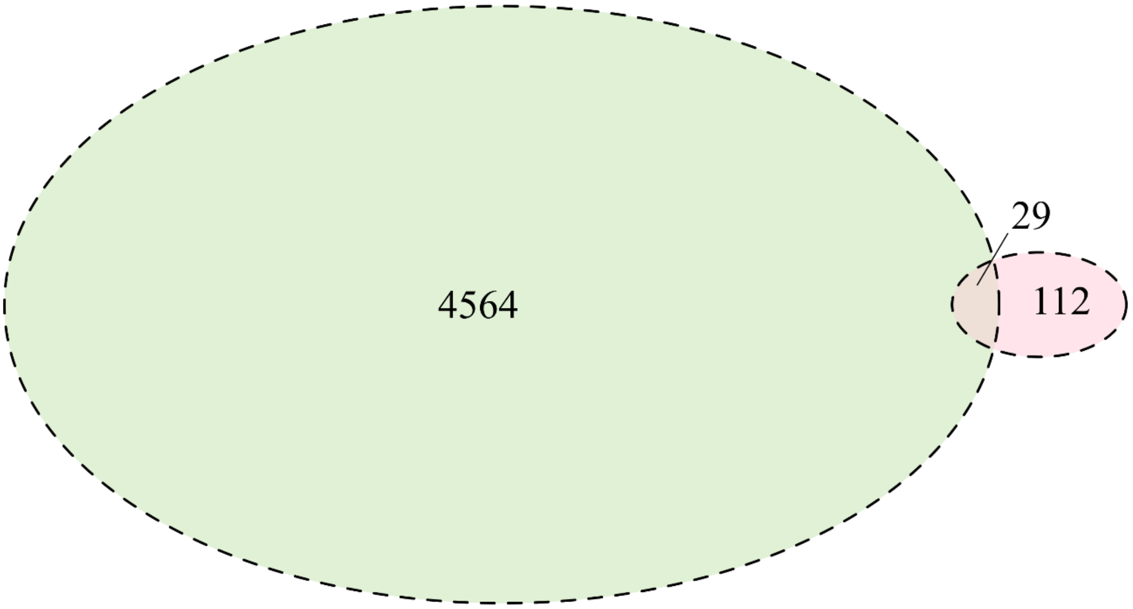


**Figure S7. Overlap of genes across datasets after therapy (T2).** Venn diagram illustrating the overlap of genes identified at T2 with genes associated with differentially methylated cytosines reported in a previous DNA methylation study by Witt et al. (green), and with genes identified in this gene expression analysis (pink). Numbers indicate the count of genes unique to or shared between the respective datasets.
